# Myo-REG: a portal for signaling interactions in muscle regeneration

**DOI:** 10.1101/711168

**Authors:** Alessandro Palma, Alberto Calderone, Andrea Cerquone Perpetuini, Federica Ferrentino, Claudia Fuoco, Cesare Gargioli, Giulio Giuliani, Marta Iannuccelli, Luana Licata, Elisa Micarelli, Serena Paoluzi, Livia Perfetto, Lucia Lisa Petrilli, Alessio Reggio, Marco Rosina, Francesca Sacco, Simone Vumbaca, Alessandro Zuccotti, Luisa Castagnoli, Gianni Cesareni

**Author notes:** Correspondence: Prof. Gianni Cesareni (G.C.).

## Abstract

Muscle regeneration is a complex process governed by the interplay between several muscle resident mononuclear cell populations. Following acute or chronic damage these cell populations are activated, communicate via cell-cell interactions and/or paracrine signals, influencing fate decisions via the activation or repression of internal signaling cascades. These are highly dynamic processes, occurring with distinct temporal and spatial kinetics. The main challenge toward a system level description of the muscle regeneration process is the integration of this plethora of inter- and intra-cellular interactions.

We integrated the information on muscle regeneration in a web portal. The scientific content annotated in this portal is organized into two information layers representing relationships between different cell types and intracellular signaling-interactions, respectively. The annotation of the pathways governing the response of each cell type to a variety of stimuli/perturbations occurring during muscle regeneration takes advantage of the information stored in the SIGNOR database. Additional curation efforts have been carried out to increase the coverage of molecular interactions underlying muscle regeneration and to annotate cell-cell interactions.

To facilitate the access to information on cell and molecular interactions in the context of muscle regeneration, we have developed Myo-REG, a web portal that captures and integrates published information on skeletal muscle regeneration.

The muscle-centered resource we provide is one of a kind in the myology field. A friendly interface allows users to explore, approximately 100 cell interactions or to analyze intracellular pathways related to muscle regeneration. Finally, we discuss how data can be extracted from this portal to support *in silico* modeling experiments.

## Introduction

Tissue homeostasis and repair after damage, are complex processes finely regulated by the interactions between different cell types exchanging signals that, in turn, stimulate intracellular signaling cascades. These orchestrated processes have been extensively studied over the past decades (Bentzinger et al. 2013; Goichberg 2016; Rojas-Ríos and González-Reyes 2014). However, the relevant information is dispersed in the bio-medical literature and scientists face the problem of designing experiments and interpreting results in the context of scattered and poorly organized prior knowledge. Although our understanding of regeneration processes is far from complete, the progress in the field would benefit from the assembly of models that take into account most of the available information. To address this issue, we conceived a web resource to organize prior knowledge about muscle regeneration processes in a structured and easily accessible format.

The adult skeletal muscle is a rather stable tissue, characterized by a relatively low cell turnover in physiological conditions (Rojas-Ríos and González-Reyes 2014). At the same time, it is remarkably dynamic and plastic and is capable of fast recovery and healing following damage caused by exercise, immobilization or chemically-induced injury (Bentzinger et al. 2013; Ceafalan, Popescu, and Hinescu 2014; Tedesco et al. 2010; Judson, Zhang, and Rossi 2013; Fu, Wang, and Hu 2015). Complete repair after damage is mediated by the regeneration process which involves the collaborative cross-talk between cells of the immune system, Fibro/Adipogenic Progenitors (FAPs) and satellite cells (SCs), among others (Bentzinger et al. 2013; Tidball 2017).

Following injury, muscle regeneration is organized into two closely interdependent phases, eventually leading to the formation of new myofibers. An initial pro-inflammatory wave triggers the recruitment of mast cells, neutrophils and monocyte/macrophage to the injury site. These cells coordinate the clearance of cellular and tissue debris, while secreting pro-inflammatory cytokines and growth factors stimulating SC proliferation and self-renewal (Bentzinger et al. 2013). At a later stage, an anti-inflammatory phase stimulates SCs to differentiate into myoblasts, which in turn fuse to form myotubes and myofibers, thus restoring the tissue architecture (Cornelison 2018).

The process is regulated in time and space and the different cell types are recruited and/or activated with distinct temporal kinetics. Myopathies and aging, interfere with this finely regulated process causing a decrease of the regeneration potential and the disruption of muscle structural organization (A Uezumi et al. 2014).

To summarize, the experimental information related to muscle regeneration is complex, diverse and sparse. Defining a strategy to combine this information in an integrated picture represents a major task. Here, we present Myo-REG, a repository of data related to muscle regeneration that was developed to meet this challenge. Myo-REG aims at capturing information on the mechanisms underlying the regeneration process of the skeletal muscle tissue. The manually-curated literature-based information is organized in a structured format and offered to the user via a friendly and intuitive interface.

Myo-REG allows users to browse a variety of data types through a single platform at https://myoreg.uniroma2.it. The stored information is annotated by manually mining the literature or by automatically capturing information from other online resources.

Myo-REG is built on two major information layers: the cell-cell interactions and the molecular interactions. Cell-cell interactions include both direct physical contacts between cells and relationships mediated by secreted signaling molecules. The molecular layer integrates physical and causal relationships that support the propagation of signals in the different cell types. The portal content can be explored starting from a cell interaction map offering an entry point for looking into both cell and signaling interactions

## Materials and Methods

### 1. Data model

Cell-cell and causal protein interactions were captured via an extensive literature search supported by text mining. To improve the consistency of the data and to make the annotations exchangeable with other resources, whenever available, we used external identifiers (IDs) for both cell and molecular entities. Since cell types are not uniquely defined and are often heterogeneous, delineating the cells that are involved in muscle regeneration required some blunt decisions from our side. This, sometimes, is the result of a compromise between alternative opinions. Cell identifiers are annotated according to the definition in the Cell Ontology section of Ontobee (Ong et al. 2017). Not all the cell types that we have considered in our model have a definition and an identifier in Ontobee. Whenever this was the case, we adopted an internal Myo-REG identifier (supplementary table 1). In addition, we used Uniprot IDs (The UniProt Consortium 2017), Chebi IDs (Hastings et al. 2016), for proteins and chemicals respectively and and SIGNOR IDs (Livia Perfetto et al. 2016) for, phenotypes and protein complexes.

For effectual usage of the stored information, interaction annotation must be based on clearly defined ontologies. This issue has been recognized since long by the molecular interaction community that, within the PSI-MI initiative, has defined ontologies that have guided the annotation of protein interaction information in Myo-REG (Sivade et al. 2018; L Perfetto et al. 2018). On the other hand, no community defined standard for cell interactions has yet been agreed upon. Thus, we developed a cell interaction ontology which is based on Gene Ontology terms (The Gene Ontology Consortium 2019; Ashburner et al. 2000) and describes the interaction mechanisms that are considered in our cell interaction model. We took into consideration both differentiation mechanisms (e.g., FAPs differentiate into adipocytes) or stimulation/inhibition mechanisms mediated either by direct cell contact or by secretion of cytokines that target a particular cell type (e.g., eosinophils secrete IL-4 to activate FAPs).

A classical logic model does not allow representing relations of the type, for instance, “TNFα inhibits FAP adipogenesis”, where adipogenesis is not a node but rather a differentiation edge between FAPs and adipocytes. Thus, in these “transition/differentiation” edges we introduced a *virtual* transition node (i.e. FAPs->adipocytes) allowing to represent that some stimuli might impact on the process.

### 2. Annotations

Cell-cell and causal protein interactions were captured by an extensive literature survey. In summary, each entity relationship stored in the database is annotated with the ID of the interacting cells, the relation mechanism, the PubMed ID of an article presenting experimental information that support the interaction and a representative sentence extracted from the same article. Although in the annotation we have retained the information about the organism where any given relationship was shown experimentally, the protein nodes in the Myo-REG signaling network are mapped to the human proteome, irrespective of the organism used to demonstrate the interaction.

Biomarkers for each cell type were selected for their relevance in the context of muscle regeneration scientific literature. This was decided after consulting relevant reviews (Bentzinger et al. 2013; Tedesco, Moyle, and Perdiguero 2017; Wosczyna and Rando 2018) describing the specific biomarkers that characterize the distinct cell types. Since biomarkers of the same cell type may differ in human and mouse, we have annotated two different biomarker tables.

### 3. Generation of Boolean models

Deriving Boolean models from causal networks implies associating to each node of the network a Boolean function stating how the node value changes according to the values of the upstream nodes. However, this operation requires establishing logic gates governed by the operators *AND*, *OR* or *NOT* to describe how two or more inputs combine to produce a single input. However, this information is not contained in causal networks and often the experimental information that is necessary to define logic gates is not available. As a consequence, the algorithm that we have implemented to translate the causal information into Boolean logic is based on the following ad hoc rules:

- The formation of a complex is associated to an ***AND*** operator (e.g. “nodeAB = nodeA *AND* nodeB”). The complex subunits must all be ‘True” for the complex to be formed.
- Inhibitions are assumed to dominate over activations. If nodeC is activated by nodeA and inhibited by nodeB, this is translated with an ***AND NOT*** operator (“nodeC = nodeA *AND NOT* nodeB).
- In establishing all the remaining updating rules, two incoming signals are combined with the ****OR**** operator (e.g. “nodeC = nodeA *OR* nodeB” if node C is activated by both nodeA and nodeB).
- When more inhibitions are present, inhibitors are combined with an ****OR**** operator (e.g. “nodeC = *NOT* (nodeA *OR* nodeB)”).

Although these assumptions often accurately describe biological interactions, they are arbitrary and may turn out to be incorrect in some cases. Whenever additional experimental information is available, the integration of activating and inactivating inputs should be evaluated critically and the corresponding logic gates edited accordingly in order to obtain a more accurate model.

### 4. Resource backbone

The resource informatics-backbone is encoded in PHP5 (http://php.net/docs.php) and JavaScript (https://www.javascript.com/) and the page styles are implemented with CSS3 (https://www.w3.org/). The database management system is PostgreSQL (https://www.postgresql.org/) that is object-relational and has a high standard compliance.

Cell-cell interaction graphs are laid out with cytoscape.js (Franz et al. 2016), a graph library written in JavaScript that allows the assembly of graphs based on entity nodes (cells and proteins) and edges (interactions among entities), including annotations for nodes and edges. Molecular interactions are displayed via the SIGNOR graph viewer (Calderone and Cesareni 2018), which allows a compartmental visualization of molecular entities as a signed directed graph.

## Results

As for many complex and dynamic biological processes, muscle regeneration can be better described by multilevel modeling (Maus, Rybacki, and Uhrmacher 2011). In this approach, interactions between cell populations in the healing tissue and activation of pathways in the different cell types denote different levels of an organizational hierarchy. Each level has its own dynamics, usually faster for molecular than for cell interactions, which are integrated by multi-level modeling to describe the behavior of the whole system.

### 1. Resource data-types and organization

The two interaction types, cell-cell and signaling interactions, are captured from the literature by manual or automatic methods (Müller et al. 2018; Orchard et al. 2012). However, being conceptually different, the two interaction types require different ontologies for annotation and data structures for organization. While hardly, any resource exists to annotate and store cell interactions, several online databases, such as KEGG (Kanehisa et al. 2017), Reactome (Fabregat et al. 2016), and SIGNOR (Livia Perfetto et al. 2016), annotate pathway-related literature information. However, they are not focused on muscle regeneration, and filtering information relevant to myologists is not straightforward.

In addition, other data types that are relevant for modeling the muscle system, such as, modulation of gene expression, are stored in generalist resources (Edgar, Domrachev, and Lash 2002; Parkinson et al. 2007), making data integration a complex task. Myo-REG aims at capturing all these information layers, in order to integrate the relevant data into a resource that can be easily explored.

The core of the Myo-REG data structure consists of two layers each describing interactions between different biological entities. The two-layered organization allows users to explore both cellular and molecular interactions. In the first layer entities are cells or secreted molecules that cells use to communicate (cytokines, growth factors and other signaling molecules). Cell interactions are defined as those relations in which either the source or the target entity is a cell, as defined in supplementary table 1. In Myo-REG cell interactions can be displayed as a static graph (diagram in the homepage) (Fig. 1, https://myoreg.uniroma2.it/) or as interactive view via a Cytoscape web application that can be accessed by clicking the “Cytoscape view” button in the home page (Franz et al. 2016). This latter representation also shows the cytokines that mediate the interactions between the cell types. In both displays, transitions (differentiations), stimulations or inhibitions are represented in different colors for a clearer interpretation. Differently from the graph view in the homepage, the Cytoscape representation is editable. Each node can be moved around for a clearer display or clicked to expand the graph to visualize the entity first neighbors. Users are also offered the possibility to remove nodes from the rendered graph. All the relationships, displayed graphically via Cytoscape, are also reported in a table format at the bottom of the page.

**Figure 1.**
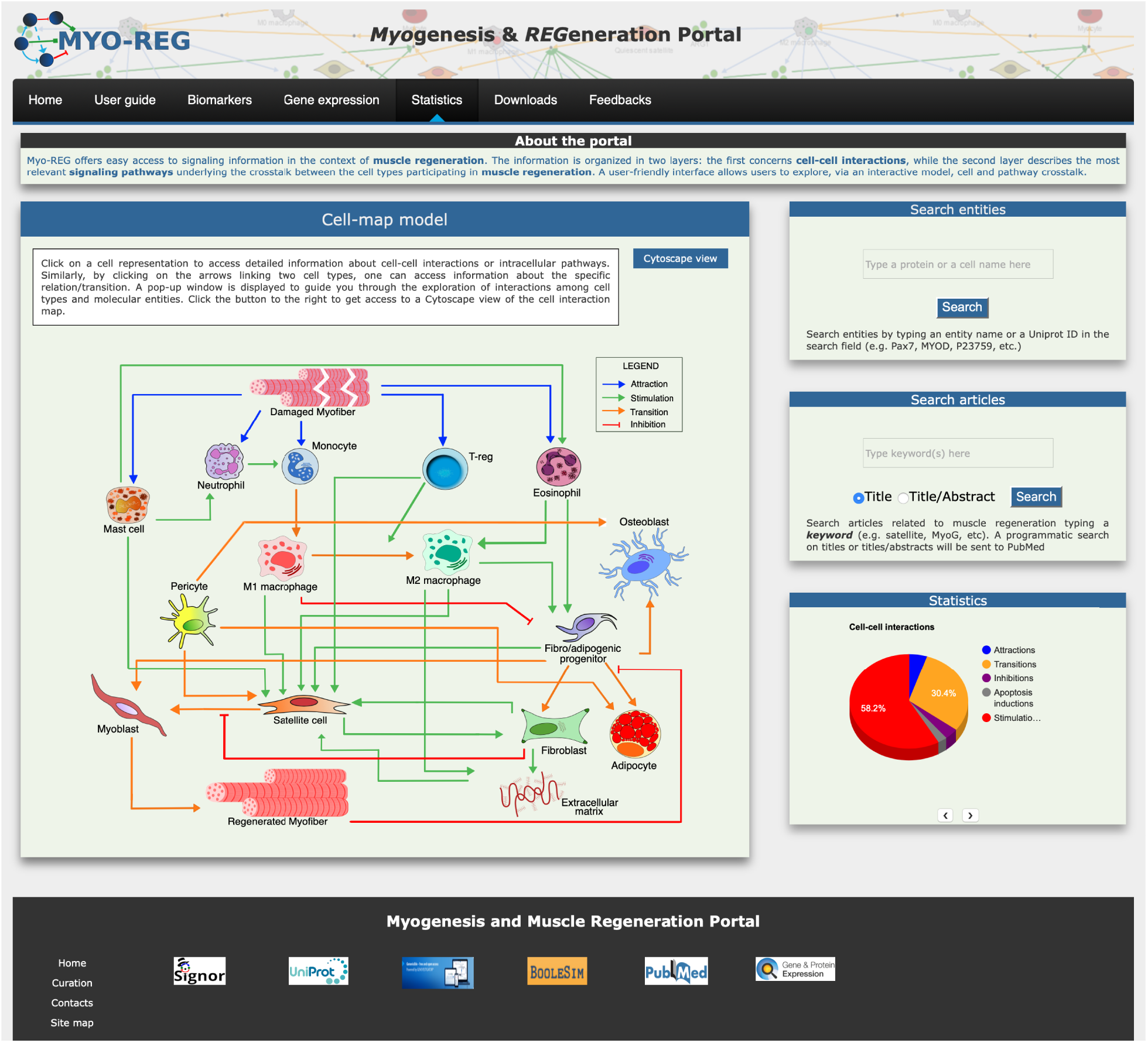
Organization of the Myo-REG home page. Myo-REG allows users to explore several aspects of muscle regeneration. The data types that are annotated in the resource include cell and signaling interactions, cell biomarkers and gene expression data. Blue arrows denote (chemo-)attractions, green arrows denote stimulations, orange arrows are transitions (mostly differentiation processes) and red t-shaped arrows are inhibitions.

The second layer annotates the molecular interactions, occurring inside the different cell types in response to external stimuli. Only relationships that are deemed relevant to muscle regeneration are considered. Differently from other signaling database, such as Reactome, that uses a reaction-based model, signaling relationships in Myo-REG are represented as causal relationships. According to this model signaling molecules are linked by edges that are directed (from modifiers to targets) and signed (activates/inactivates).

Finally, in Myo-REG causal interactions are organized into pathways. The cell interactions in the muscle, either by direct contact or via secretion of cytokines, impact on the activation of pathways in target cells, thus changing their phenotypes. Pathways are abstractions linking a subset of signaling molecules by a chain of causal relations. For each cell type in Myo-REG, we manually curated pathways defining a set of causal interactions that explain the cell response to molecular signals, in the context of muscle regeneration. Pathways are shown as signed directed graphs that link an input signal to an output phenotype. The definition of the molecular entities that should be included in a pathway is subjective. Most pathways in Myo-REG include as few as 10-20 entities (mini-pathways) and initiate with an environmental signal to terminate with a phenotype. The compartmental representation of the pathway graphical display provides a user-friendly way to organize the signaling cascade in distinct cellular spaces: extracellular, plasma membrane, cytoplasm and nucleus. To date, Myo-REG stores a collection of 76 pathways that can be displayed and explored in each of the distinct cell-type pages (supplementary table 2).

### 2. Home page

The resource home page is organized to offer easy access to the annotated information and to search tools. It contains a navigation toolbar and four frames displaying i) an interactive cell-map model, (ii) a form to search the database for the presence of annotated information on cell or protein entities, (iii)a form to search the literature and finally iv) a fourth frame displaying some database statistics (Figure 1).

The main entry point to access the stored information is a cell-map model linking the different cell types (Figure 1 and 2A). Both nodes and edges in the cell interaction graph are actionable.

**Figure 2.**
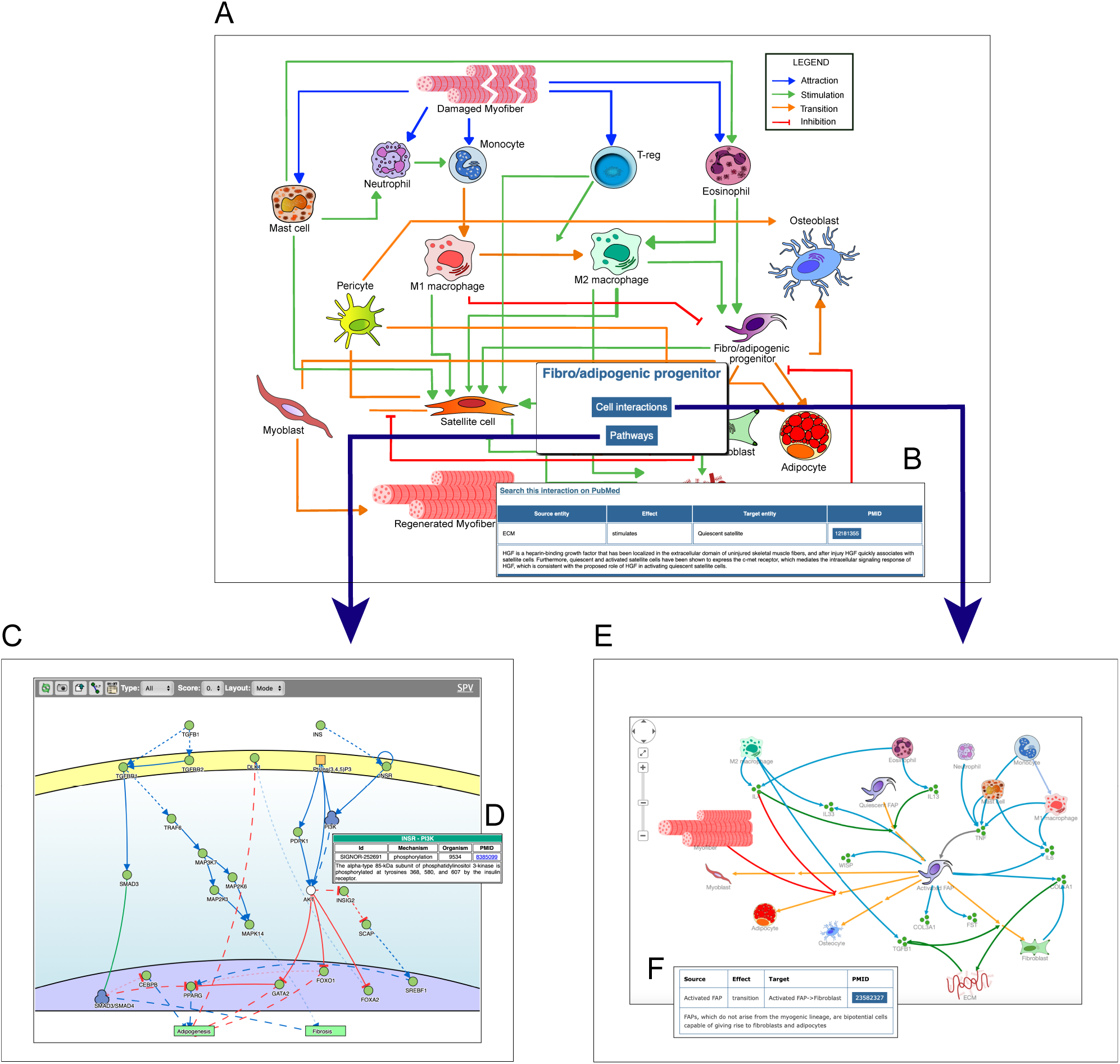
Navigating the Myo-REG resource. A) In the homepage a network of the cell interactions occurring in muscle regeneration is the main entry point to the information annotated in Myo-REG. By clicking a cell icon, a pop-up window is displayed offering the possibility to select either the “Cell interactions” or the “Pathways” tabs. (C) By selecting the pathway tab the user can choose to display a network representation of the pathways that are relevant for that cell type. (D) The annotation of an interaction (e.g., INSR-PI3K) displayed after clicking the edge on the graph. (E) The cell-cell interactions graph displayed for a selected cell type (FAP). (F) The annotation of an interaction displayed after clicking on an edge of the graph. Edge colors represent distinct interaction types according to the legend displayed on the top-left corner of the graph. Icons with three green circles represent cytokines participating in the process of muscle regeneration.

The user can select any cell type of interest and, by clicking the “cell interaction button” in the pop-up window he/she can choose to explore outgoing and incoming edges that modify the cell behavior. This information is returned as a Cytoscape view (Fig. 2 E) showing first and second neighbors of the selected cell. Edges are colored according to the type of interaction and annotated with: i) the gene names of the entities that take part in that interaction, ii) the effect on the target entity iii) the PubMed ID of the article reporting experimental evidence supporting the relation and (iv) a representative short sentence describing the specific relationship (Fig. 2 F).

Alternatively, users can explore intracellular pathways that modulate the differentiation potential of the selected cell or its ability to secrete cytokines that affect the activation of other cell types (Fig. 2 C). For each cell type, the page allows to display the pathways that are deemed relevant to explain the function of that cell type. As said, pathways include 10-20 signaling molecules (mini-pathways). However, users can assemble larger pathways by choosing to display more than one mini-pathway (via check-boxes) in order to explore pathway cross-talks.

In addition, by clicking on a graph edge, a pop-up window appears showing information about the selected cell relationship including a hyperlink to the PubMed page of the article reporting the experimental observations that support the interaction (Fig. 2B). Finally, the hyperlink “search this interaction in PubMed” in the pop-up window launches a PubMed query looking for manuscripts containing the names of the two interacting cells in association with the “muscle” term in the title or abstract (Fig. 2 B).

### 3. Search tools and statistics

Two additional frames in the home-page right side give access to search tools. The “search entities” tool allows exploring the role that a gene product or a cell type plays in muscle regeneration. Briefly by typing the protein/gene name or protein ID (Uniprot ID) in the search entity-form the tool returns a table that reports the protein matches in the database. Users can choose, via check boxes, to display either the signaling partners of the query protein or the Myo-REG mini-pathways the protein is involved in. Alternatively, if the query entity is a cell type, the result page displays a table allowing to choose whether to display a Cytoscape view of the cells interacting with the query cell or the signaling pathways that are associated to that cell in Myo-REG. A second search form allows to retrieve articles containing any keyword in connection with the term “muscle” in an article title or abstract. A programmatic query is sent to PubMed to retrieve articles whose content may be related to the query term in the context of muscle regeneration. The search results are reported as a table presenting the title, PMID, author list, publication title, publication year and journal name.

### 4. The navigation toolbar

Additional data and tools can be accessed via the navigation toolbar (Figure 3).

**Figure 3.**
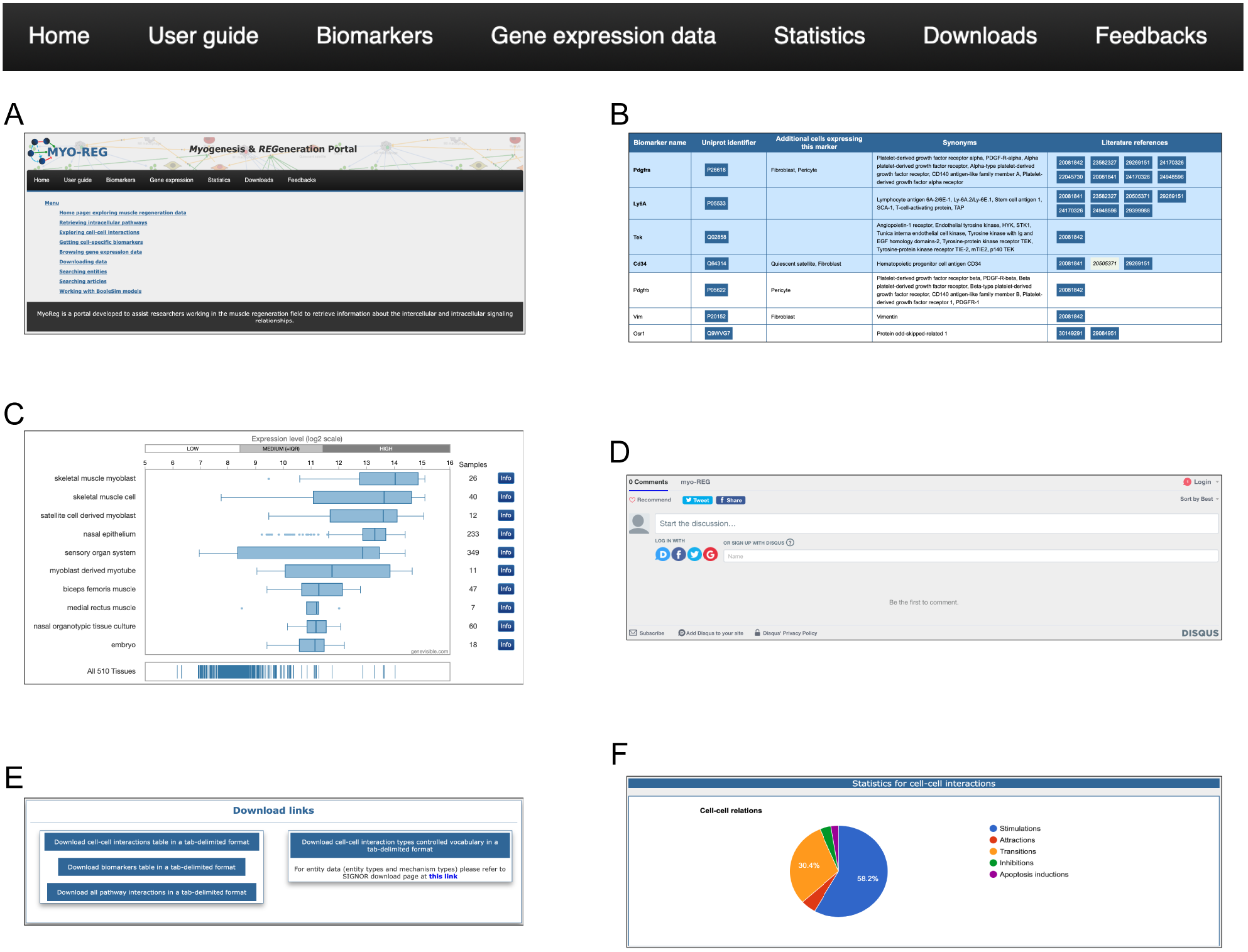
The navigation toolbar. The navigation toolbar in the home page allows users to explore additional website content, including a short user-guide (A), a list of cell type specific biomarkers (B), gene expression data (C), a feedback page (D) where users can suggest additions, modifications or corrections, a download page (E) for downloading information for local use and finally a statistics page (F)

A first tab, “gene expression”, links to the Genevisble web service (Hruz et al. 2008) (Fig. 3 E). Genevisible is a curated resource that integrates public gene expression data from both microarray and RNA-seq experiments. By entering a query gene name and by selecting either *Homo sapiens* or *Mus musculus* the resource returns the list of 10 tissues, cell lines or cancers with the highest expression of the query gene. The results are displayed as boxplots.

Muscle resident cells are classified into different cell types according to the presence of antigens that are differently expressed in the different cell types. In Myo-REG these antigens are dubbed “Biomarkers”. Biomarkers can be accessed by clicking the “Biomarkers” tab in the navigation bar (Fig. 3 B). This action leads to a search form that allows to select the organism (*H. sapiens* or *M. musculus*) and, from a drop-down menu, the cell type of interest. The search results are displayed in a table format where each biomarker is reported with its gene name and synonyms, the UniProt identifier, a list of additional cell types expressing that biomarker and reference to the literature supporting its expression in the selected cell type.

To assist the resource exploration, we have also compiled a user-guide (Fig. 3 D) describing in more detail and illustrating with self-explanatory screenshots the main operations that a Myo-REG user could consider. All the data annotated in Myo-REG can be downloaded from Myo-REG for local use via the “download” tab. Finally, the “statistics” tab gives access to a variety of analyses of the database content.

### 5. Executable Boolean models

To facilitate hypothesis generation, we implemented a script to convert logic into Boolean models for any pathway of interest curated in Myo-REG. The models are then displayed via a customized BooleSim application (Bock et al. 2014). BooleSim is a simple network simulator, which allows to create, edit and simulate networks based on Boolean logic. The automatically generated models can be modified, for instance by editing Boolean rules, thereby allowing to introduce into the model additional information. The generated models can be used to perform *in silico* simulations under different conditions. To illustrate the potential of this tool we will assemble and analyze a model of FAP differentiation when stimulated by insulin and TGFβ.

FAPs are muscle resident interstitial cells of mesenchymal origin (Joe et al. 2010; Akiyoshi Uezumi et al. 2010; Stumm et al. 2018). When activated, they transiently support satellite cells proliferation through the establishment of several paracrine signals (Joe et al. 2010; Murphy et al. 2011). In pathological conditions, such as in Duchenne muscular dystrophy, FAPs can differentiate into fibroblasts or adipocytes, contributing to the deposition of fat tissue and to the formation of scar infiltrations (Akiyoshi Uezumi et al. 2011). When treated ex vivo with insulin (INS) FAPs differentiate into adipocytes (Joe et al. 2010), while the treatment with TGFβ triggers their fibrogenic differentiation (Akiyoshi Uezumi et al. 2010).

Myo-REG stores curated pathways describing FAP response to INS and TGFβ. In Fig. 4 we have merged the two pathways and automatically converted the resulting causal network into an executable Boolean network.

**Figure 4.**
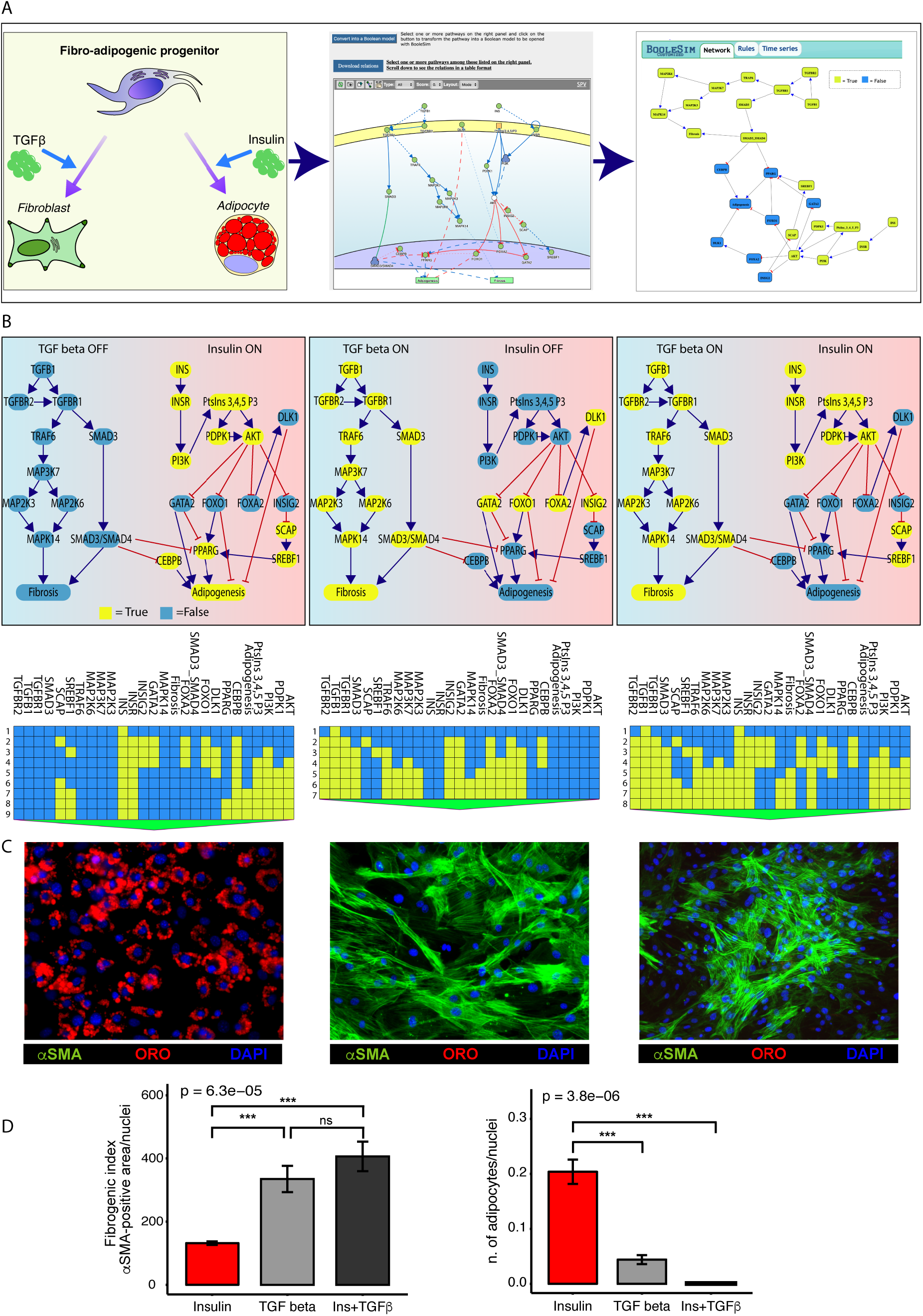
Conversion of the TGFb and INS pathways into an executable Boolean network. (A) The ability of FAPs to differentiate into either adipocytes or fibroblasts (left panel) is represented in Myo-REG as a union of two selected pathways: INSR and TGFb (middle panel). Pathways are automatically converted into a Boolean network and input into a customized Boolesim application (right panel). (B) The panel depicts the system stationary phase in conditions in which the FAPs are stimulated with insulin (left panel), TGF beta (middle panel) or both (right panel). The network evolution is represented as a heatmap below each simulation. Yellow means active (True) while blue is inactive (False) (C) Representative immunofluorescence (20x magnification) of FAPs from wild type mice stimulated to differentiate in the presence of insulin (left), TGF beta (middle) or both stimuli (right). Adipocytes (red) and fibroblasts (green) were stained with oil Red O (ORO) and with an anti-α-SMA antibody respectively. Nuclei (blue) were stained with Hoechst 33342. (D) Left chart: quantification of the percentage of green area with insulin treatment as reference group. Right chart: quantification of the number of adipocytes normalized over the number of nuclei per field (insulin treatment as reference group).

The network encompasses 27 nodes linked by negative (inhibitions) or positive (activations) relationships. The nodes can have a value of either “*True*” or “*False*” (yellow and blue in Fig. 4 respectively). During the simulation steps the value of each node is updated according to the values of the upstream nodes and to the rules defined in the Methods section. Only two nodes (INS and TGFb) have no incoming edge and represent the input of the system, while “Adipogenesis” and “Fibrosis” are output nodes (nodes without outgoing edges) and are used as read-outs. In the simulation in Figure 4 all nodes have an initial value set to *False*, except for the input nodes that define the experimental conditions. The values of the readouts when the system is stabilized, however, only depend on the initial values of the input nodes, irrespective of the initial values of the remaining nodes.

As shown in the left panel of figure 4 B, when the value of the INS input node is set to “*True*” (yellow) and the TGFb to “*False*” (blue) after a few simulation steps the system reaches a steady state were the adipogenesis node is *True* and fibrogenesis is *False*. Conversely when TGFb is set to *True* and INS is *False* the model predicts that at steady state fibrogenesis is *True* (Fig. 4 B middle panel). The times series data displayed below each panel show the evolution of the system starting from the initial condition. This model prediction is coherent with experimental observations. Interestingly, the model predicts that, when FAPs are given both the INS and TGFb stimuli, the antiadipogenic TGFb signal wins over the pro-adipogenic insulin signal and drives the system into fibrogenesis (Fig. 4 B right panel). Also this prediction was verified experimentally (Figure 4 C-D and supplementary file 3).

## Discussion

Myo-REG is a web portal designed to facilitate access to organized information in the context of muscle regeneration. The user-friendly interface and the variety of annotated experimental information are intended to help myologists to integrate the different data-types that are relevant for their work.

Myo-REG is a multi-scale, manually-annotated, resource that is one in a kind, since it provides information on how intracellular molecular events affect cell phenotypes and cell-communication during muscle regeneration.

Although we aimed at high coverage and accuracy, we are aware that our effort is likely to have missed some relevant piece of information while some relationships might have been annotated incorrectly. We are committed to continue curating this resource to improve accuracy and coverage by including new data, as published. This can better be achieved if colleague myologists, that find this resource useful, interact with us and send feedback and suggestions for improvements and corrections. A tab in the navigation bar opens a “feedback” form that allows users to comment on data annotation and suggest inclusion of additional information or corrections.

## Acknowledgements

Not applicable

## Authors’ contributions

A.P., L.C. and G.C. conceptualized the project. All authors contributed to data curation. In particular A.P. curated cell-cell interactions, pathways and biomarkers, L.L.P. and C.F. curated pathways and biomarkers, A.C.P, F.F., C.G., G.G, M.I., L.L., A.R., M.R., F.S., S.V., S.P., L. P. and A.Z. curated pathways, L.C. and G.C curated cell-cell interactions, pathways and biomarkers. A.P. developed the database, web site and scripting. A.C. developed the pathway graph viewer and contributed to the website conceptualization E.M. implemented the Boolean network transformation script. A.P., A.R., M.R. and G.C. conceptualized the wet lab experiment. A.P. performed the cell culture, treatments, immunofluorescences, image acquisition and statistical analyses. A.P. wrote the original draft. A.P., C.G., L.C. and G.C. wrote and edited the manuscript. A.P. and G.C. prepared the figures. All authors read and approved the final manuscript.

## Availability of data and materials

**Project home page**: https://myoreg.uniroma2.it

**Operating system(s):**Platform independent

**Programming language:**Html5 for the website backbone, Css3 for styles, Php5, R and JavaScript for the scripting part.

**Other requirements:**optimized for Mozilla Firefox and Safari browsers

The data used and/or analyzed during the current study are available from the corresponding author on reasonable request

