## Supplementary file 3 for "Myo-REG: a portal for signaling interactions in muscle regeneration"

### FAPs isolation, culture conditions and treatments

*Mdx* FAPs were isolated as Cd31^-^/Cd45^-^/α7-integrin^-^/Sca1^+^, according to Reggio et al. 2019 (1).

FAPs were resuspended in growth medium (GM), composed of 20% FBS, 10 mM Hepes, 1 mM sodium pyruvate, 100 U/ml P/S in high glucose DMEM GlutaMAX™, and seeded at the cell density of 4 × 10^4^ cell/well. Two days after plating, the GM was fully refreshed and cells cultured with differentiation media. The adipogenic differentiation was induced by incubating FAPs with the adipocyte differentiation medium (ADM) composed of GM + 1 µg/ml human recombinant insulin, 0.5 mM 3-isobutyl-1-methylxanthine (IBMX) and 1 μM dexamethasone. Two days after treatment cell were cultured for three additional days in adipocyte maintenance medium (AMM: GM + 1 µg/ml insulin). The fibrogenic differentiation was induced by incubating FAPs with 5 ng/ml of human recombinant transforming growth factor-β (TGFb) in 20% FBS for two days, followed by three days in 20% FBS medium. The adipogenic/fibrogenic cotreatment was performed by incubating FAPs in ADM containing TGFb for two days, followed by three days in AMM.

### Immunofluorescence

Cells were fixed in 2% PFA for 10 minutes at RT and permeabilized in 0.5% Triton X-100, followed by saturation of non-specific binding sites with blocking solution (10% FBS, 0.1% Triton X-100 in PBS 11X) for 1 hour at RT. Primary antibody (α-SMA) was diluted in blocking solution and incubated for 1 hour at RT. Cells were washed four times with 0.1% Triton X-100 in PBS 1X and incubated for 30 minutes at RT with anti-mouse secondary antibody diluted in blocking solution. Cells were washed four times with 0.1% Triton X-100 in PBS 1X, stained with Oil Red O staining and counterstained for 5 minutes at RT with 1 mg/ml Hoechst 33342 in 0.1% Triton X-100 in PBS1X.

### Oil Red O staining

Oil Red O (Sigma Aldrich, catalog O0625) solution was prepared according to the manufacturer’s recommendations. Fixed cells were washed twice in PBS 1X and incubated for 10 minutes with the staining solution (Oil Red O with ultra-pure water in a 3:2 ratio). Stained cells were washed three times with 1X PBS and counterstained using Hoechst 33342. Oil Red O stained cells were acquired via fluorescence microscopy.

### Images acquisition and analysis

The immunofluorescences were automatically acquired via LEICA fluorescent microscope (DMI6000B). Twenty images per well were acquired at 20x magnification.

### Cell differentiation measurements and statistical analysis

Adipogenic differentiation of FAPs was estimated using ImageJ by counting the number of adipocytes (ORO-stained) normalized over the total number of nuclei per field. 20 fields were analyzed for each treatment at 20X magnification.

Fibrogenic differentiation was assessed using ImageJ by estimating the SMA-stained area for each field (expressed in pixels). 20 fields were analyzed for each treatment at 20X magnification.

Data are reported as mean ± SEM. Statistical significance was measured using the unpaired Student’s t-test, with significance values reported as follows: *p < 0.05; **p < 0.01; ***p < 0.001. Statistical analyses were performed using R software (2) “ggplot2” package (3).

### Antibodies used in this study

| **Antibody** | **Vendor** | **Catalog number** | **Immunofluorescence** |
| --- | --- | --- | --- |
| α-Smooth Muscle Actin (α-SMA) (1A4) | Sigma Aldrich | A5228 | 1:300 |
| Anti-mouse Alexa Fluor® 488 | Molecular probe | A11001 | 1:200 |
